# OrgaMeas: A pipeline that integrates all the processes of organelle image analysis

**DOI:** 10.1101/2024.04.15.589634

**Authors:** Taiki Baba, Akimi Inoue, Susumu Tanimura, Kohsuke Takeda

## Abstract

Although image analysis has emerged as a key technology in the study of organelle dynamics, the commonly used image-processing methods, such as threshold-based segmentation and manual setting of regions of interests (ROIs), are error-prone and laborious. Here, we present a highly accurate high-throughput image analysis pipeline called OrgaMeas for measuring the morphology and dynamics of organelles. This pipeline mainly consists of two deep learning-based tools: OrgaSegNet and DIC2Cells. OrgaSegNet quantifies many aspects of different organelles by precisely segmenting them. To further process the segmented data at a single-cell level, DIC2Cells automates ROI settings through accurate segmentation of individual cells in differential interference contrast (DIC) images. This pipeline was designed to be low cost and require less coding, to provide an easy-to-use platform. Thus, we believe that OrgaMeas has potential to be readily applied to basic biomedical research, and hopefully to other practical uses such as drug discovery.

## Introduction

Organelles are intracellular compartments within eukaryotic cells, and are represented by nuclei, endoplasmic reticulum, and mitochondria. They perform unique functions such as protein synthesis, protein modification, and energy production to support cellular function and homeostasis. Therefore, organelle dysfunction can be reflected in a wide variety of pathological conditions (Saminathan et al., 2021). Because of their multiple physiological and pathophysiological roles, the interest in analyzing organelles from multifaceted perspectives has been increasing in recent years. Accordingly, microscopy techniques and organelle labeling methods have been developed and widely used, making it possible to accurately observe the intracellular dynamics of organelles (Chin et al., 2021). Ideally, such image analysis technology should encompass quantitative measurement techniques, and the development and widespread use of free image analysis software, such as ImageJ/Fiji, have provided many researchers with a user-friendly platform for quantitatively analyzing biomedical images (Schneider et al., 2012). Furthermore, ImageJ/Fiji allows researchers to readily build macros that can automate a series of image processing steps and handle large amounts of imaging data. Thus, ImageJ/Fiji is currently most commonly used for organelle image analysis.

Organelle image analysis is generally performed following the three steps of segmentation, setting of regions of interest (ROIs), and quantification. Using image analysis software, such as ImageJ/Fiji, organelle images are segmented using a thresholding method available within the software. In such a method, image data are binarily divided into foreground pixels representing organelles and background pixels representing non-organelle regions based on an intensity threshold determined by the researchers. Prior to quantifying morphological metrics of the organelles of each cell, researchers usually manually set ROIs to segment individual cells. However, there are some serious problems with the commonly used image analysis procedures (Miura and Nørrelykke, 2021). One is that the currently used thresholding methods are not necessarily accurate. Both the low signal/noise ratio of images and the presence of a broad range of fluorescence signals frequently leads researchers to fail to detect positive signals or to mistakenly pick up background signals. Another problem is that the manual setting of ROIs is a laborious task for researchers, and therefore the throughput of such procedures can be quite low. More seriously, both thresholding methods and ROI settings are influenced by researcher subjectivity, and are therefore prone to bias. Thus, to gain reliability, reproducibility, and throughput in the image analysis of organells, a new technology that can solve these problems is required.

In the past few years, artificial intelligence technologies based on deep learning, a family of machine learning techniques, have been applied to basic biomedical research (Moen et al., 2019). In the cell biology field, deep learning has enabled significant improvements in image analysis, including in the segmentation of intracellular structures (Gallusser et al., 2023), prediction of organelle-specific fluorescence signals from unlabeled microscopy images (Trizna et al., 2023), denoising (Laine et al., 2021), and super-resolution image reconstruction (Xu et al., 2023). U-Net is a popular deep learning technique that is specifically used to develop segmentation tools for biomedical images (Ronneberger et al., 2015). MitoSegNet, a representative deep learning tool that was recently developed using U-Net, has achieved highly accurate image segmentation of mitochondria (Fischer et al., 2020). Deep learning has also been exploited to establish segmentation tasks for other organelles, such as the nuclei (Heckenbach et al., 2022) and lysosomes (Morone et al., 2020), thereby demonstrating that this emerging technology can contribute a great deal to the advancement of organelle image analysis. However, the segmentation tools so far developed do not exert their full performance in the final quantification of morphological metrics of organelles unless each tool is coupled with an automated, accurate high-throughput tool for ROI setting, which is currently not readily available or accessible. Moreover, most segmentation tools were developed analyze a particular organelle of interest, and useful open tools that cover multiple organelles have not been developed. Therefore, the cell biology research community is currently in need of a versatile pipeline integrating tools for segmentation, ROI setting, and quantification in the image analysis of different organelles.

To provide such a pipeline, we developed OrgaMeas in this study. OrgaMeas consists mainly of two image processing tools, OrgaSegNet and DIC2Cells, which were developed with the deep learning techniques U-Net and pix2pixHD, respectively. OrgaSegNet precisely segments different organelles, and DIC2Cells automates ROI settings based on differential interference contrast (DIC) images, enabling high-throughput and highly-accurate analysis of organelle features at the single-cell levels. Moreover, we demonstrate the scope of application of OrgaMeas by showing that it can effectively and accurately quantify the morphological metrics of different organelles and changes to them under several experimental conditions.

## Results

### Optimization for organelle imaging

To acquire images suitable for precise, quantitative analyses, we sought to determine the experimental conditions required for labeling organelles in cultured cells. To this end, we focused on the mitochondrion as a representative organelle and HeLa cells as representative cultured cells because they are widely used in a variety of biological research. Using this organelles and cell type, we compared the following three techniques: immunofluorescence of fixed cells, live imaging of chemically labeled cells, and live imaging of genetically labeled cells (**Fig. S1**). Immunofluorescence that detected the outer mitochondrial protein Tom20 in fixed cells yielded mitochondrial images with lower resolution than was achieved by either live imaging of cells labeled with the fluorescent compound MitoTracker Green FM, which is incorporated into mitochondria, or live imaging of cells expressing the transgene encoding Mito-DsRed, a mitochondrial matrix-targeted fluorescence protein. This lower resolution with immunofluorescence may be because the fixation step affected the integrity of the mitochondrial membrane and/or proteins. When the two live imaging techniques were compared, live imaging of chemically labeled cells appeared to be better than live imaging of genetically labeled cells because the latter tended to cause mitochondria to be aberrantly swollen, which may be caused by overexpression of exogenous mitochondrial proteins. Therefore, we adopted live imaging of chemically labeled cells as the optimal method for organelle imaging.

### Development of DIC2Cells, a deep learning-based tool for cell segmentation in DIC images

In the image analysis of organelles, certain features of them are generally quantified at the single-cell level by setting single-cell ROIs. To visualize cellular regions and annotate each cell, dyes that stain the cell membrane or cytoskeleton are often used. However, it is preferable to avoid the labeling of cells with such dyes because it can raise concerns of cytotoxicity and damage to organelles. Thus, we sought to exploit the power of deep learning to develop a tool that enables the setting of single-cell ROIs without any labeling of cells (**Fig. 1**).

**Figure 1.**
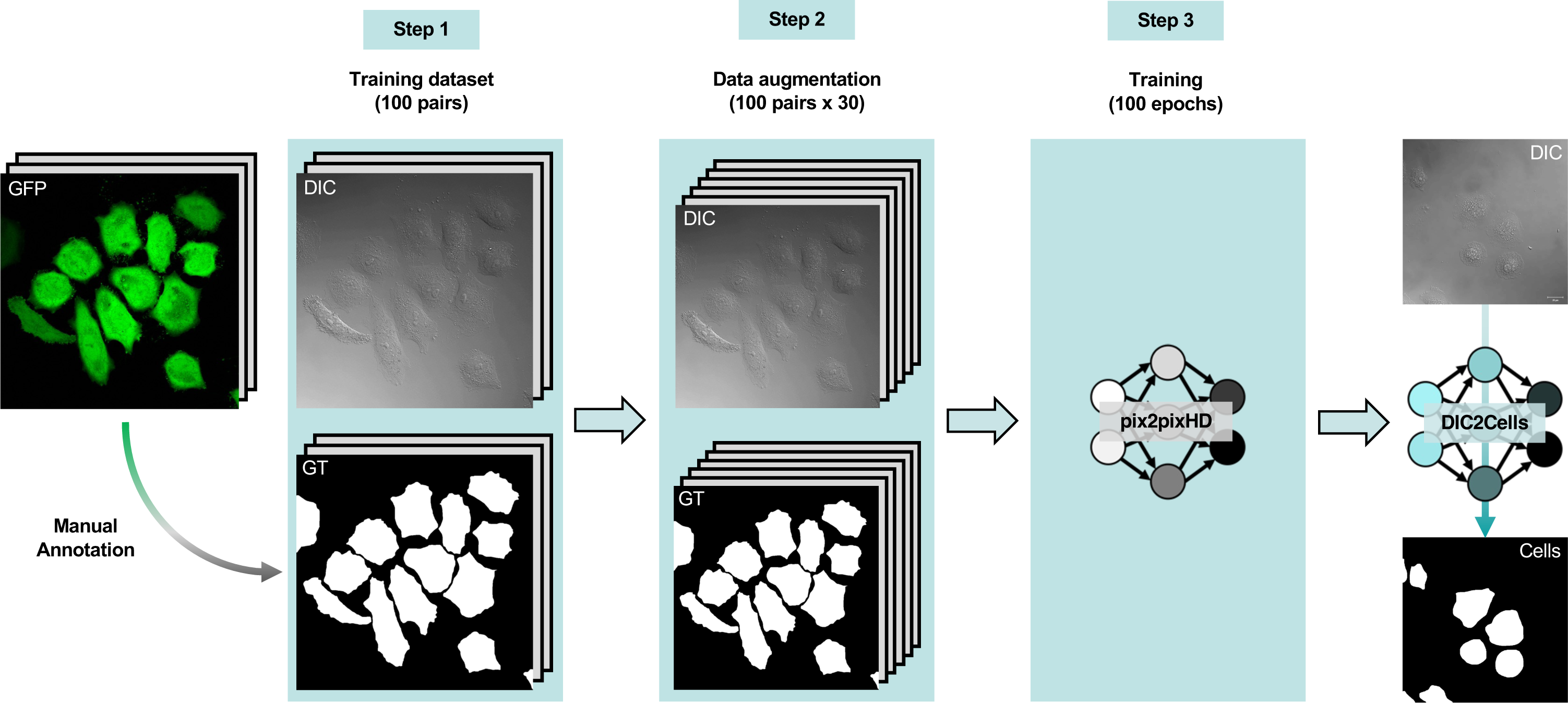
Development of DIC2Cells, a deep learning-based tool for cell segmentation in DIC images. Workflow of the development of DIC2Cells through to the training of pix2pixHD. Differential interference contrast (DIC) and green fluorescent protein (GFP) images were acquired through live-cell imaging of HeLa cells stably expressing GFP. The training dataset consisted of 100 pairs of DIC and ground truth (GT) images, the latter being manually generated by segmenting individual cells according to their GFP signals (**Step 1**). The volume of the training dataset was augmented 30 times using augmentation operations (**Step 2**). Finally, pix2pixHD was trained with the training dataset using optimized training hyperparameters (**Step 3**).

We adopted DIC microscopy as the acquisition method for cell images, because it is often compatible with live cell imaging for the acquisition of organelle images and yields images with higher contrast and less halo than those obtained by other light microscopy techniques such as bright field and phase-contrast microscopy (Piper and Piper, 2014). To segment individual cells in DIC images, we used pix2pixHD, which is a deep learning framework for high-resolution image-to-image translation (Wang et al., 2018), and was recently applied to the segmentation of electron microscopy images (Jerez et al., 2021). To generate training data for pix2pixHD, we first acquired green fluorescent protein (GFP) and DIC images through live-cell imaging of HeLa cells stably expressing GFP. Using the GFP images, we created ground truth (GT) images by carefully annotating and binarizing individual cells according to their GFP signals, which corresponded well to the cell regions, finally generating a training dataset consisting of 100 pairs of DIC-GT images (**Fig. 1**, Step 1). Because the accuracy of deep learning models generally increases with the quantity, as well as the quality, of training data (Mahajan et al., 2018), we used data augmentation methods to increase the volume of the training dataset by 30 times (**Fig. 1**, Step 2**; Table S1**). Finally, this training dataset was used to train pix2pixHD, and 10 models, which we designated as the DIC2Cells models, were obtained by saving the trained data at every 10 epochs until 100 epochs (**Fig. 1**, Step 3).

### Validation of DIC2Cells

We next validated the cell segmentation accuracy of DIC2Cells following the workflow indicated in **Fig. 2A**. In the same manner as that used to prepare the training dataset, another dataset consisting of 30 pairs of DIC-GT images that were not involved in the training step, was generated, and its volume was augmented 10 times (**Fig. 2A**, Steps 1 and 2**; Table S2**). The 10 models of DIC2Cells (10-epoch to 100-epoch) were then used to predict cell segmentation images (Cells) from the DIC images in this validation dataset (**Fig. 2A**, Step 3). To validate the cell segmentation accuracy, we first compared the predicted images with their respective GT images (which were manually segmented from the DIC images) and calculated the Dice coefficient, a frequently used metric for evaluating pixel-wise similarity between two images (**Fig 2A**, Step 4). A representative image comparing a GT image and Cells image is shown in **Fig. 2B**. The 10 models consistently achieved Dice coefficient values as high as 0.96 to 0.97 (**Fig. 2C**). We next evaluated how accurately the DIC2Cells models distinguished between neighboring cells and found that the accuracy appeared to be inconsistent among the models (**Fig. S2**). To address this point, we calculated the cell count accuracy (CCA) and found that it varied from 0.874 to 0.974 independently of the number of epochs (**Fig. 2D**). On the basis of these data, we finally adopted the most accurate 40-epoch model as the final DIC2Cells model. We also developed an ImageJ/Fiji macro called “Single Cell Creator” that automatically generates single-cell organelle images from the outputs of DIC2Cells, thereby accomplishing a seamless workflow for the automated ROI setting (**Fig. 5A**).

**Figure 2.**
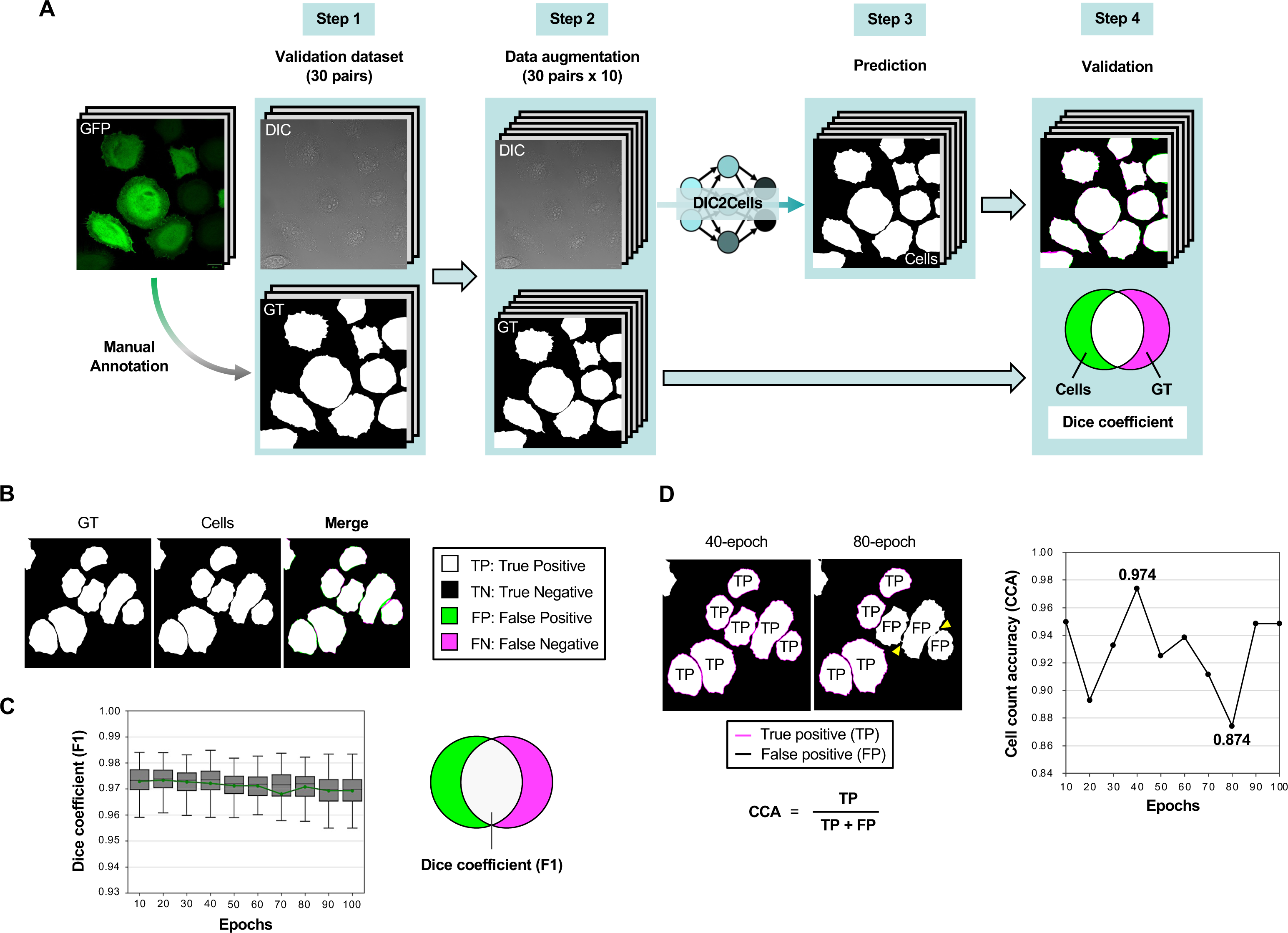
Validation of the accuracy of DIC2Cells in cell segmentation. **(A)** Workflow to validate the cell-segmentation accuracy of DIC2Cells. Differential interference contrast (DIC) and GFP images were acquired through live-cell imaging of HeLa cells stably expressing GFP. The validation dataset consisting of 30 pairs of DIC images and ground truth (GT) images was prepared (**Step 1**). The volume of the validation dataset was augmented 10 times using the augmentation operations (**Step 2**). Each of the DIC2Cells models obtained by saving the trained data at every 10 epochs was used to predicte cell-segmented images (Cells) from the DIC images in the validation dataset (**Step 3**). Finally, we compared the Cells images with their respective GT images and calculated the Dice coefficient (**Step 4**). **(B)** Comparison between the GT and Cells images, where the true positive segmentation, the true negative segmentation, the false positive segmentation (negative in GT; positive in Cells), and the false negative segmentation (positive in GT; negative in Cells) are shown in white, black, yellow-green, and magenta, respectively. **(C)** The Dice coefficient values (F1) achieved by the 10 DIC2Cells model. **(D)** The cell count accuracies (CCA) of the 10 DIC2Cells models (Right graph). Left panels show representative images generated by the 40-epoch and 80-epoch models, which were the most and least accurate models, respectively. Yellow arrowheads indicate incomplete distinction between neighboring cells, which resulted in a false cell area in which three cells were counted as one. True positive (TP) indicates the number of cells, which were successfully segmented compared with the GT images, whereas false positive (FP) shows the number of cells, which were incompletely distinguished from neighboring cells.

### MitoSegNet effectively segments certain organelles other than mitochondria

We next focused on the segmentation of organelles in live-cell imaging data. Although the efficiency and accuracy of organelle segmentation have much improved with the recent emergence of deep learning technologies, most of the segmentation tools developed so far are aimed at the analysis of a particular organelle of interest, and useful open tools that cover multiple organelles have not been developed. To develop a segmentation tool that addresses this issue, we first evaluated MitoSegNet, a deep learning-based tool dedicated to the precise segmentation of mitochondria (Fischer et al., 2020). As has been reported, we found that MitoSegNet accurately segmented not only typical tubular mitochondria but also fragmented mitochondria with a vesicle-like or rod-like appearance (**Fig. 3A**). This raised the possibility that MitoSegNet could be applied to segmentation of other vesicle-like organelles. Indeed, we demonstrated that MitoSegNet accurately segmented lysosomes without further training, including those faintly labeled with fluorescent dyes in the raw data (**Fig. 3B**). MitoSegNet also segmented lipid droplets, which had a similar round appearance to lysosomes but were larger (**Fig. 3C**). However, MitoSegNet failed to segment nuclei (which are also round organelles but have a much larger size) and yielded irregular mosaic patterns within nuclear regions, probably because each nucleus was heterogeneously stained with the fluorescent dyes (**Fig. 3D**). These results suggest that MitoSegNet can be applied to segmentation of not only mitochondria, but also other organelles with a similar form, as long as they can be represented by a homogeneously-labeled area.

**Figure 3.**
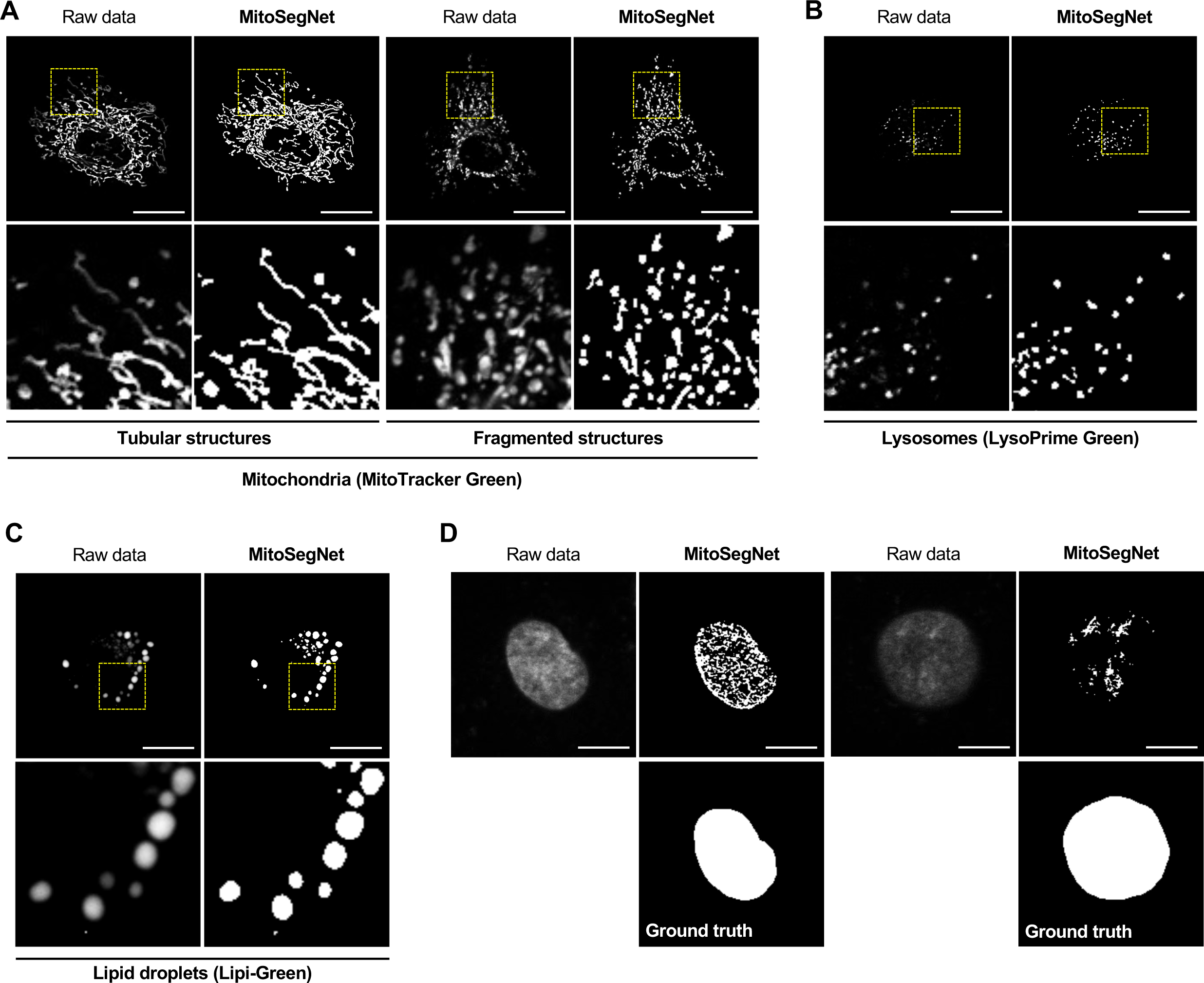
MitoSegNet effectively segments organelles other than mitochondria. **(A-C)** Representative results of organelle segmentation by MitoSegNet. Images of mitochondria, lysosomes, and lipid droplets (Raw data) were acquired using live-cell imaging of HeLa cells stained with MitoTracker Green **(A)**, LysoPrime Green **(B)**, and Lipi-Green **(C)**, respectively. Scale bar, 10 µm. The lower panels are magnified images of the regions surrounded by yellow dotted lines in the respective upper panels. **(D)** Representative results of nuclear segmentation by MitoSegNet. The images of nuclei were obtained by live-cell imaging of HeLa cells stained with Hoechst 33342. Scale bar, 10 µm. Ground truth images show ideal segmentation results for the respective raw nuclear data.

### Development of NucSegNet, a deep learning-based tool for segmentation of the nuclei

Thus far, we have described our use of the pre-trained MitoSegNet model, which comes in a ready-to-use “basic mode” that can segment mitochondria without further training. In addition to this mode, Fischer et al. provide an “advanced mode” that allows researchers to create a custom segmentation model by means of U-Net, a deep learning framework for the segmentation of biomedical images (Ronneberger et al., 2015). Using this latter mode, we developed a deep learning tool for the segmentation of nuclei, which we called NucSegNet. To begin with, we acquired images of nuclei by live-cell imaging of HeLa cells stained with Hoechst 33342 and created their respective GT images by manually annotating and binarizing the individual nuclei, generating a dataset that consisted of 20 pairs of nuclei-GT images (**Fig. 4A**, Step 1). Each image in the training dataset was then split into four tiles, and the volume of the dataset was increased by 40 times with data augmentation (**Fig. 4A**, Step 2**; Table S3**). Finally, 80% of the dataset was used to train U-Net, and the remaining 20% was used to validate the accuracy of the resulting models in the segmentation of nuclei (**Fig. 4**, Step 3). During the training, the loss function (which is an indicator of how well the training is going) was monitored at each epoch, and the training was completed at 20 epochs, where both the training loss and validation loss were minimal (**Fig. 4B**). Consistent with this training procedure, the segmentation accuracy quantified by the Dice coefficient was highest at 20 epochs with both the training and validation data (**Fig. 4C**). In fact, the 20-epoch model successfully segmented nuclei with a wide range of labeling intensities, without being affected by heterogeneous labeling within each nuclear region (**Fig. 4D**). Thus, we finally adopted the 20-epoch model as the final NucSegNet model.

**Figure 4.**
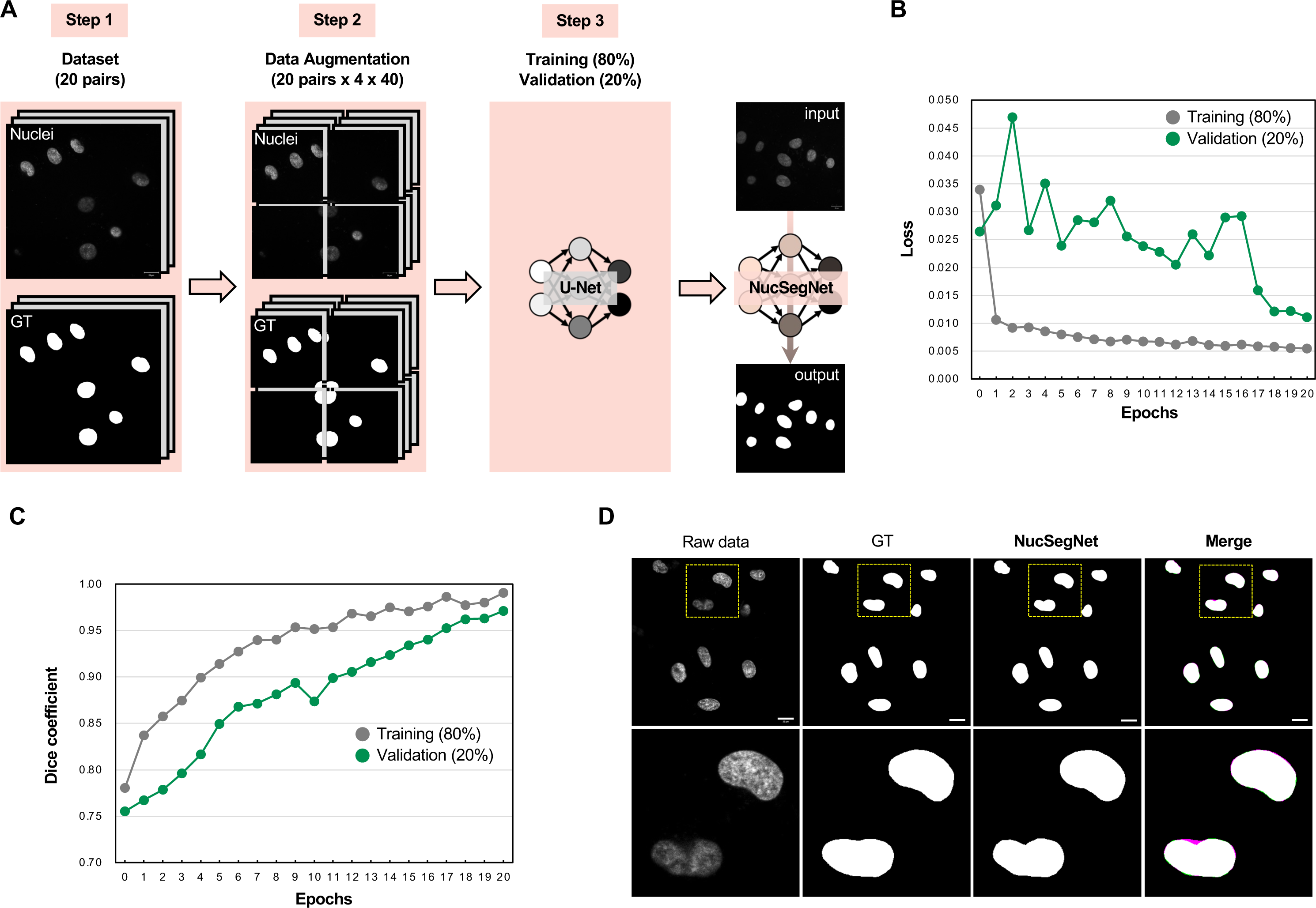
Development of NucSegNet, a deep learning-based tool for segmentation of the nuclei. **(A)** Workflow of the development of NucSegNet through to the training of U-Net in the “advanced mode” of MitoSegNet. Images of nuclei were acquired by live-cell imaging of HeLa cells stained with Hoechst 33342. A dataset consisting of 20 pairs of nuclear images and their respective ground truth (GT) images was prepared (**Step 1**). Each image in the training dataset was split into four tiles, and then the volume of the dataset was increased 40 times with data augmentation (**Step 2**). Finally, U-Net was trained with 80% of the training dataset and optimized hyperparameters, and the Dice coefficient was calculated for the other 20% of data reserved for validation (**Step 3**). **(B)** Loss functions for training (80% of the dataset) and for validation (the remaining 20%) are shown for each epoch. **(C)** The averages of training and validation Dice coefficients calculated with 80% of the dataset and the remaining 20%, respectively. **(D)** Representative images of nuclei stained with Hoechst 33342 (Raw data), the ground truth results (GT), the output of NucSegNet, and the overlap between the GT and output of NucSegNet (Merge). Scale bar, 20 µm. In the segmented image, the true positive segmentation, true negative segmentation, false positive segmentation (negative in GT; positive in NucSegNet), and false negative segmentation (positive in GT; negative in NucSegNet) are shown in white, black, yellow-green and magenta, respectively. The lower panels are magnified images of the region surrounded by yellow dotted lines in the respective upper panels.

### OrgaMeas effectively and accurately quantifies many aspects of different organelles

Taking advantage of our findings that MitoSegNet is applicable to certain organelles other than mitochondria and our development of the new deep learning-based tool NucSegNet, we combined these two tools to construct a more versatile organelle segmentation tool that we named OrgaSegNet. To make the most of OrgaSegNet, we also developed an original ImageJ/Fiji macro called Organelle Analyzer that automatically quantifies various organelle parameters, such as the number, morphology, and intracellular distribution, using segmented images obtained by OrgaSegNet (**Fig. 5B**). Furthermore, we finally integrated the OrgaSegNet–Organelle Analyzer unit for segmenting organelles with the DIC2Cells–Single Cell Creator unit for automated ROI setting, thereby completing an organelle image analysis pipeline called OrgaMeas that enables high-throughput highly-accurate analysis of organelle features at the single-cell level (**Fig. 5**).

**Figure 5.**
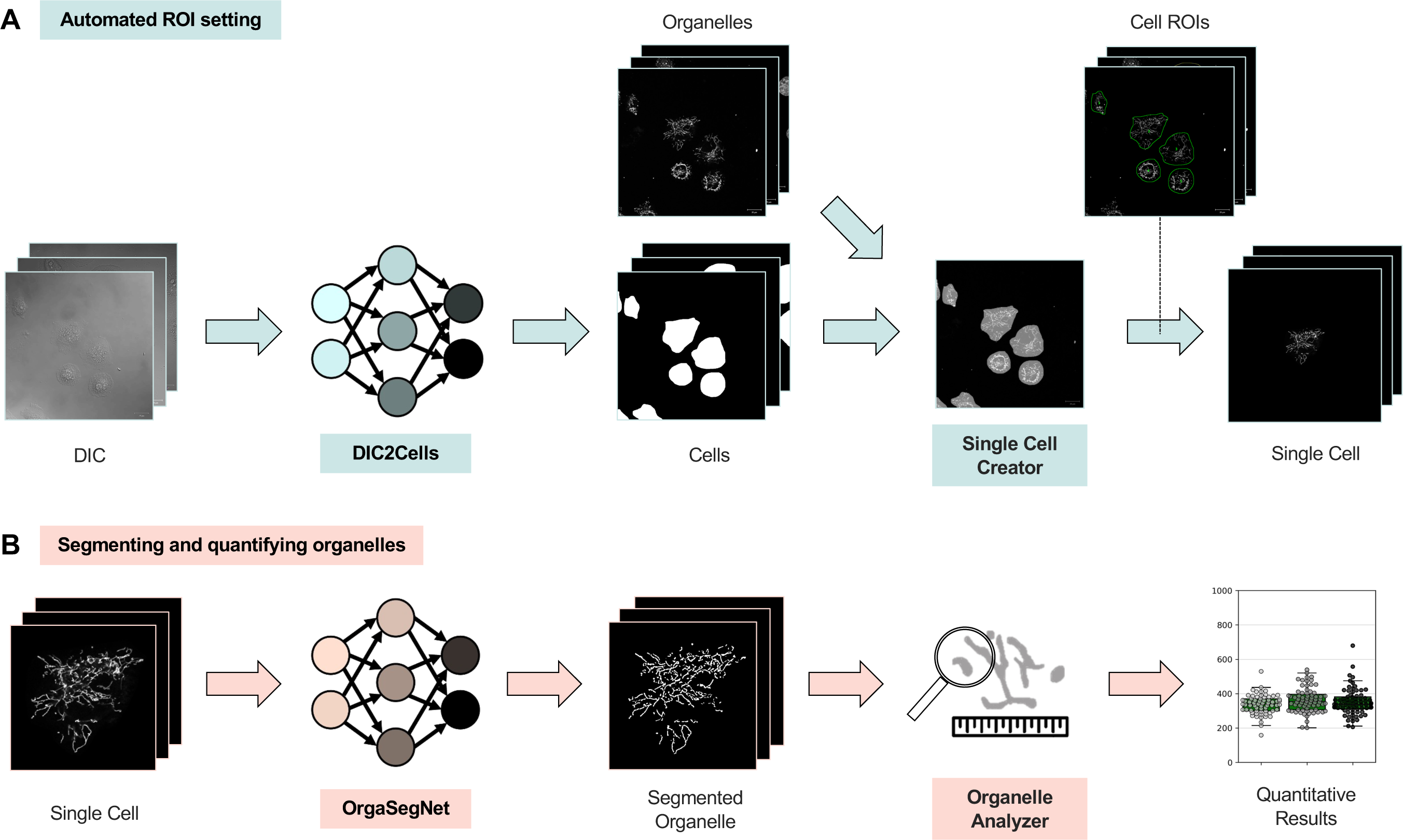
Overview and workflow of OrgaMeas. **(A)** The DIC2Cells–Single Cell Creator unit for automated ROI setting. DIC2Cells and Single Cell Creator, an Image J/Fiji macro-based tool, provide a seamless workflow for the automated creation of ROIs on DIC images. **(B)** The OrgaSegNet–Organelle Analyzer unit for segmenting and quantifying organelles. OrgaSegNet and Organelle Analyzer, another Image J/Fiji macro-based tool, automatically quantify various organelle parameters, such as the number, morphology, and intracellular distribution.

To assess the scope of application of OrgaMeas, we applied it to the analyses of mitochondria, lysosomes, and nuclei in their steady state and under some experimental conditions. We first evaluated images of mitochondria and pharmacologically induced changes in morphology in HeLa cells stained with MitoTracker Green FM. As reported in a number of studies (Tanaka and Youle, 2008; Ishihara et al., 2003), the mitochondrial uncoupler CCCP induced fragmentation of mitochondria, whereas the inhibitor of mitochondrial fission Mdivi-1 induced elongation of mitochondria (**Fig. 6A**). OrgaMeas was applied to these images and used to quantify the number and morphological metrics of mitochondria per cell for more than 100 cells per experimental group, without any laborious manual work. As shown in **Fig. 6B**, the quantitative analysis showed that CCCP induced an increase in the number of mitochondria and a decrease in the mitochondrial area, the length of the major axis, and the number of branches, all of which are hallmarks of mitochondrial fragmentation. In contrast, Mdivi-1 induced opposite changes that represent elongation of mitochondria.

**Figure 6.**
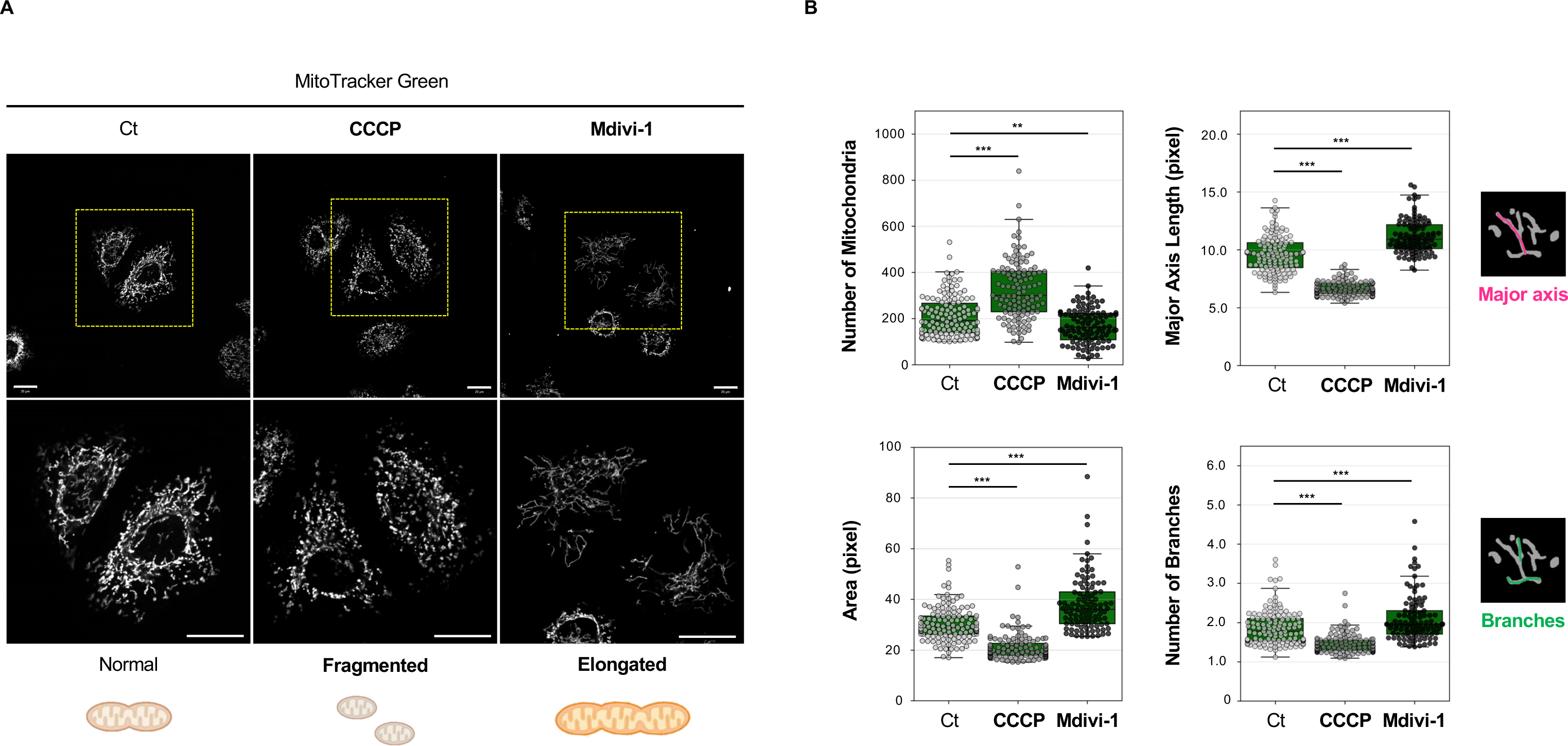
OrgaMeas effectively quantifies well-known morphological changes in mitochondria. **(A)** Representative images of HeLa cells treated with 10 µM CCCP or 50 µM Mdivi-1 for 1 h in the presence of MitoTracker Green. Scale bar, 20 µm. The lower panels are magnified images of the regions surrounded by yellow dotted lines in the respective upper panels. Representative data from three independent experiments are shown. **(B)** Quantification of the number and morphological characteristics of mitochondria per cell using OrgaMeas. More than 100 cells were analyzed per experimental group. Data are presented as individual means. **p < 0.01, ***p < 0.001, Dunnett’s multiple comparison test (Ct vs. CCCP or Mdivi-1). Representative data from three independent experiments are shown.

Next, we focused on lysosomes and changes in their morphology and activation state, the latter of which depends on the extent of acidification of their lumen. We deprived HeLa cells of serum and amino acids to create acidic conditions to activate lysosomes, and further treated the cells with the protease inhibitors pepstatin A and E-64d to inhibit protein degradation within lysosomes. Total lysosomes and activated lysosomes were stained with the pH-insensitive probe LysoPrime Green and the pH-sensitive probe LysoTracker Red, respectively (**Fig. 7A**). The fluorescent images obtained were then analyzed using OrgaMeas. The total lysosomal area and the average diameter of individual lysosomes both increased in response to starvation, suggesting that the size of individual lysosomes increased upon lysosomal activation (**Fig. 7B**). Intriguingly, treatment of the cells with the protease inhibitors further increased both of the parameters, suggesting that the activated lysosomes enlarged further. This may be because unprocessed substrate proteins accumulated in the activated lysosomes upon the inhibition of lysosomal proteases. As expected, quantitative analysis of the LysoTracker Red signal by OrgaMeas revealed that those cells that possessed lysosomes with high LysoTracker Red signal strongly increased in number in response to starvation, irrespective of the presence or absence of the protease inhibitors (**Fig. 7C**).

**Figure 7.**
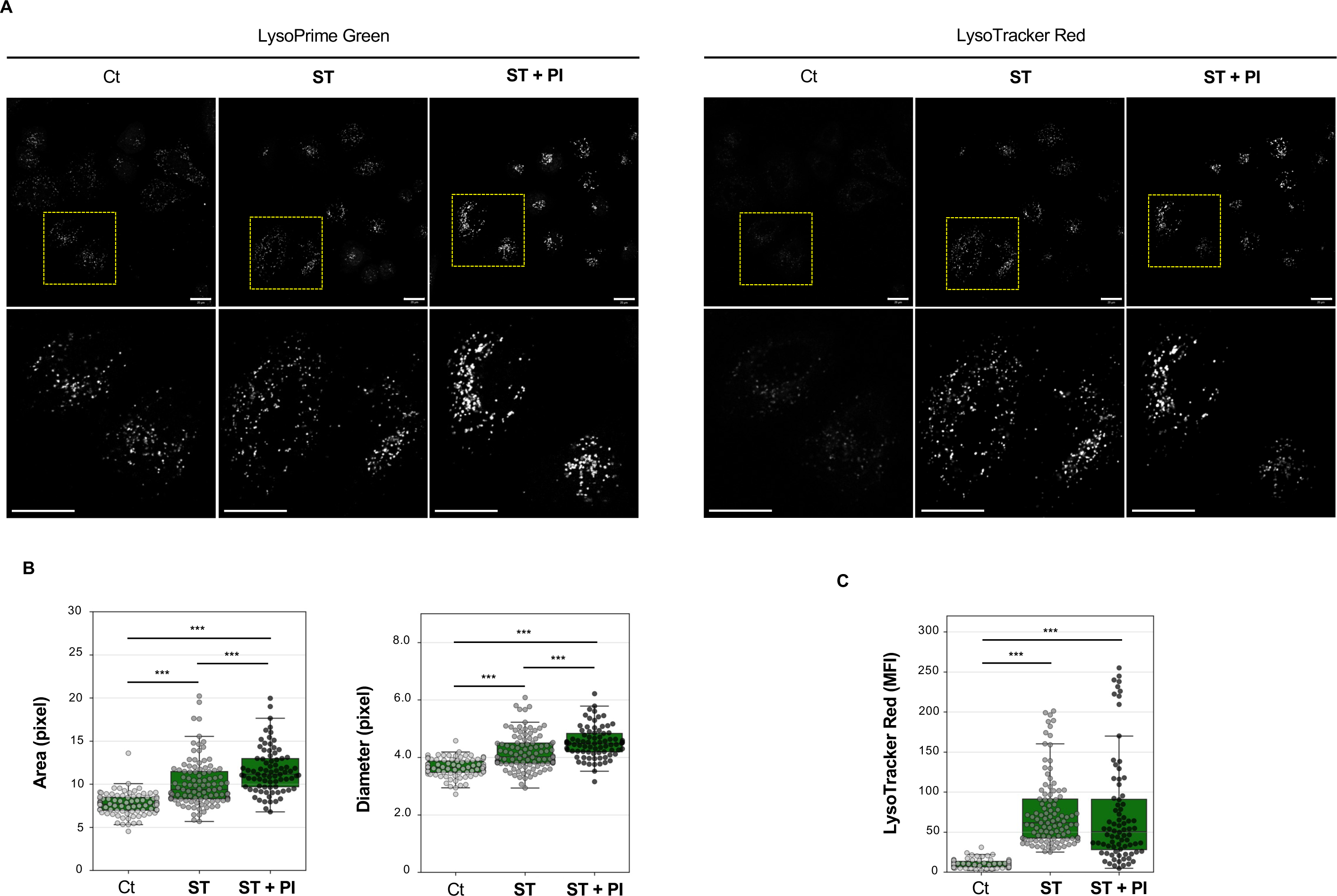
OrgaMeas quantitatively detects the changes in morphology and the activation state of lysosomes. **(A)** Representative images of HeLa cells deprived of amino acids and FBS (ST: starvation) in the presence or absence of protease inhibitors (PI), 10 μg/mL pepstatin A, and 10 μg/mL E-64d, for 1 h. Total lysosomes and activated lysosomes were stained with the pH-insensitive probe LysoPrime Green (left six panels) and the pH-sensitive probe LysoTracker Red (right six panels), respectively. Scale bar, 20 µm. Lower panels are magnified images of the regions surrounded by yellow dotted lines in the respective upper panels. Representative data from three independent experiments are shown. **(B, C)** Quantification of the lysosomal area and average diameter of individual lysosomes per cell (B) and the intensity of LysoTracker Red signal per cell (C) using OrgaMeas. More than 100 cells were analyzed per experimental group. Data are presented as individual means. ***p < 0.001, Dunnett’s multiple comparison test (Ct vs ST or ST + PI) in (B), Tukey’s multiple comparison test (Ct vs ST, Ct vs ST + PI, ST vs ST + PI) in (C). Representative data from three independent experiments are shown.

Finally, we evaluated the nuclei and their morphological changes using OrgaMeas. For this, we used cytoskeleton modifying agents to induce morphological abnormalities in the nuclei because actin filaments localize not only in the cytosol but also in the nucleus, where they play a role in maintaining the integrity of the nuclear structure (Davidson and Cadot, 2021; Gieni and Hendzel, 2008). We found that nuclear shape was obviously distorted when HeLa cells were treated with latrunculin A, an inhibitor of actin polymerization (**Fig. 8A**). Interestingly, nocodazole, an agent that interferes with the polymerization of microtubules, similarly induced abnormal nuclear shapes. To quantitatively evaluate these aberrant shapes, we obtained segmented nuclear data from OrgaMeas and used them to calculate solidity, which indicates the degree of convexity and concavity (**Fig. 8B**), and circularity, which indicates the degree of similarity to a perfect circle (**Fig. 8C**). Consistent with the imaging data, application of either latrunculin A or nocodazole decreased both the parameters, quantitatively indicating that the disturbance of cytoskeletal polymers induced the morphological complexity of the nuclei. Together, these results demonstrated that OrgaMeas can detect and accurately quantify many aspects of different organelles.

**Figure 8.**
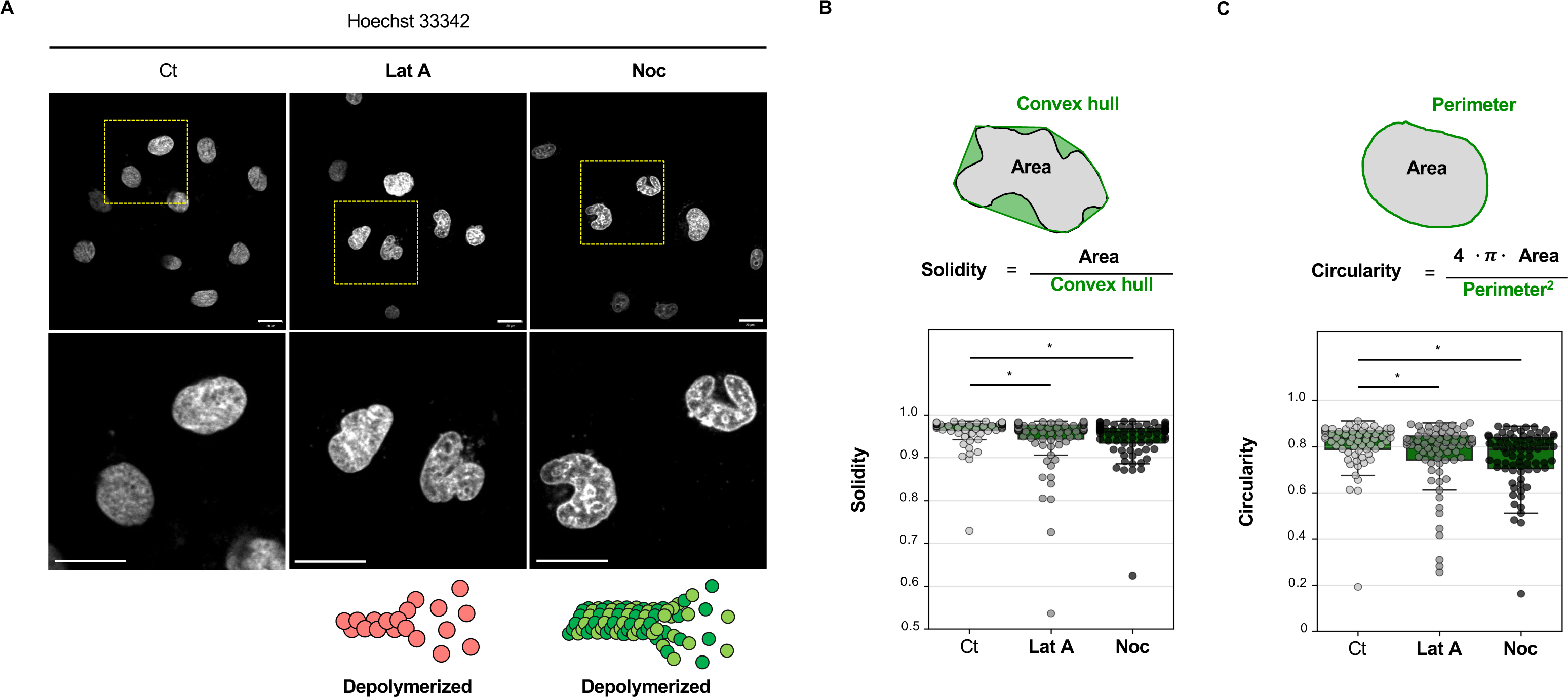
OrgaMeas quantitatively detects changes in nuclear shape. **(A)** Representative images of HeLa cells treated with either 0.1 µM latrunculin A (Lat A) or 10 µM nocodazole (Noc) for 30 min in the presence of Hoechst 33342. Scale bar, 20 μm. Lower panels are magnified images of the regions surrounded by yellow dotted lines in the respective upper panels. Representative data from three independent experiments are shown. **(B, C)** Solidity (B) and circularity (C) of the nuclei per cell calculated from the segmented data obtained using OrgaMeas. More than 100 cells were analyzed per experimental group. Data are presented as individual means. **p < 0.05, **p < 0.01, Dunnett’s multiple comparison test (Ct vs Lat A or Noc). Representative data from three independent experiments are shown.

## Discussion

With the recent developments in microscopic and organelle labeling techniques, image analysis for the assessment of organelle dynamics, such as changes in morphology and spatial distribution, has been a central approach in the study of cell biology. In addition, open image analysis software such as ImageJ/Fiji has allowed researchers to analyze large quantities of images. However, the methods commonly used in such software are error-prone and laborious, and few alternative tools to solve these problems are available. In this study, we present an automated image analysis pipeline called OrgaMeas, which enables effective and accurate quantification of organelle dynamics at a single-cell resolution.

Prior to developing this pipeline, we optimized the image acquisition methods for quantitative organelle analysis (**Fig. S1**). We found that when considering the selectivity and sensitivity of labeling, cellular toxicity, and the time and effort required, live imaging of chemically labeled cells is the best way for accurate analysis of organelles. Live cell imaging facilitates easy visualization of cellular regions using optical microscopy, without the requirement to label cells with fluorescent proteins or dyes, which may induce cytotoxicity or damages to organelles, but it is generally difficult to obtain images with a high enough signal-to-noise ratio to accurately segment individual cells. Among the various available light microscopy techniques, DIC microscopy appeared to be the most promising because it is generally compatible with live cell imaging for acquisition of organelle images, and yields images with higher contrast and less halo than can be obtained by any other light microscopy techniques, such as bright field or phase-contrast microscopy (Piper and Piper, 2014).

The past few years have seen the development of cell segmentation tools for optical microscopy that employ the power of artificial intelligence methods (Mukherjee et al., 2023; Hilsenbeck et al., 2017; Pachitariu and Stringer, 2022). Regarding the application of such techniques to DIC microcopy images, Lux and Matula reported success in cell segmentation of DIC images of HeLa cells, but unfortunately did not provide the source code or a tool (Lux and Matula, 2019). This situation motivated us to develop our own cell-segmentation tool for DIC images and release it as an open tool. To begin with, we focused on pix2pix, a deep learning technique for image-to-image translation (Isola et al., 2017). Pix2pix adopts U-Net as the neural network architecture, a network commonly used in biomedical image segmentation (Ronneberger et al., 2015), and the learning process of pix2pix employs a conditional adversarial generative network (cGAN), which enables the generation of ideal images as similar as possible to the GT images (Goodfellow et al., 2014). In this study, we employed pix2pixHD, an improved version of pix2pix that can deal with high-resolution images (Wang et al., 2018), and finally developed DIC2Cells. Together with Single Cell Creator, our original ImageJ/Fiji macro, we accomplished a seamless workflow for the automated creation of ROIs on DIC images (**Fig. 5**).

Although the recent development of deep learning technologies has greatly improved the efficiency and accuracy of organelle segmentation (Morone et al., 2020; Heckenbach et al., 2022), most segmentation tools developed to date are limited to the analysis of a particular organelle of interest, and useful open tools covering multiple organelles have not been developed. In the present study, we focused on MitoSegNet, a highly accurate mitochondria-specific segmentation tool (Fischer et al., 2020). Unexpectedly, we found that MitoSegNet can be applied not only to mitochondria but also to other organelles that have morphological similarities to mitochondria, such as lysosomes and lipid droplets (**Fig. 3**). However, because we failed to segment nuclei using MitoSegNet, we developed our own nuclear segmentation tool, NucSegNet, by employing the “advanced mode” of MitoSegNet (**Fig. 4**). We then combined these two tools to form a more versatile tool, OrgaSegNet, for one-pot quantification of different organelles, thereby paving the way to develop tools that cover a wider range of organelles. Recently, many studies have found that extensive communication through membrane contact between various types of intracellular membrane compartments is essential for cellular, tissue, and organismal function and homeostasis (Prinz et al., 2020). OrgaMeas may be helpful for analyzing such inter-organelle contacts and their significance in organelle and cellular functions.

Although many tools for biomedical image analysis have been reported, such as those for cell segmentation, ROI setting, and organelle segmentation, most of these were developed as stand-alone tools for application to a single process, and therefore the research community is demanding analysis pipelines that seamlessly cover the entire image analysis process. Although such pipelines already exist, they were developed using conventional image processing methods that are available within ImageJ/Fiji and are prone to errors (Garcia-Pardo et al., 2021; Klickstein et al., 2020). To address this issue, we took advantage of the power of deep learning and developed OrgaMeas as an organelle image analysis pipeline with high accuracy, high throughput, and low bias (**Fig. 5**). In OrgaMeas, the imaging data used for analysis are limited to DIC images because the pipeline uses DIC2Cells to perform cell segmentation. However, deep learning tools can be easily finetuned to perform different and/or additional tasks by training them with less data than is required for untrained models (Iman et al., 2023). If necessary, users can re-train and fix DIC2Cells to adapt it to other images such as bright field images and phase contrast images. This would further expand the versatility of OrgaMeas.

Although deep learning techniques have been increasingly applied in the life sciences in recent years (Moen et al., 2019), most of the tools developed are currently not widely used in the research community. This may be because many researchers lack the knowledge and skills of data science, and therefore developers should design tools with a user-friendly platform that does not require any coding or a high-spec computer. Taking this situation into account, we developed DIC2Cells for implementation in Google Colaboratory, which is widely used within the data science community for developing deep learning projects (Carneiro et al., 2018). Furthermore, we provide easy-to-understand instructions for OrgaMeas, including for DIC2Cells (https://github.com/MitoTAIKI/DIC2Cells-from-OrgaMeas), OrgaSegNet (https://github.com/MitoTAIKI/NucSegNet-from-OrgaMeas), and the ImageJ/Fiji macros (https://github.com/MitoTAIKI/IJ-macros-from-OrgaMeas), on Github. We hope that OrgaMeas will be widely used by researchers, including those who are not familiar with data science, and that beyond its uses in basic biomedical research it will also be applied to a variety of practical uses such as drug discovery.

## Materials and methods

### Reagents

CCCP, pepstatin A, and E-64d were purchased from Fujifilm Wako Chemicals (Osaka, Japan). Mdivi-1 was purchased from Tocris Bioscience (Bristol, UK). Latrunculin A and nocodazole were purchased from Cayman Chemical (Ann Arbor, MI) and Sigma Aldrich (St. Louis, MO), respectively.

### Cell culture

HeLa cells stably expressing GFP (GFP-HeLa cells) were established previously (Tanimura et al., 2011). HeLa cells and GFP-HeLa cells were cultured in Dulbecco’s modified Eagle’s medium (D-MEM) (high glucose; Fujifilm Wako Chemicals) containing 8% fetal bovine serum (FBS), 100 U/ml penicillin G, and 0.1 mg/ml streptomycin under a 5% CO_2_ atmosphere at 37°C.

### Live-cell imaging

#### Mitochondria staining

Cells were grown on glass-bottom dishes (12-mm Glass Base Dish; IWAKI, Shizuoka, Japan), treated with appropriate reagents, and simultaneously stained with 150 nM MitoTracker Green FM (Invitrogen, Waltham, MA) for 1 hour (h). After washout, cells were cultured in phenol red-free D-MEM (high glucose without L-glutamate; Fujifilm Wako Chemicals) containing 8% FBS, 4 mM L-glutamate (Sigma Aldrich), 100 U/ml penicillin G, and 0.1 mg/ml streptomycin under a 5% CO_2_ atmosphere at 37°C, and were then imaged by confocal microscopy.

#### Lysosome staining

Cells were grown on glass-bottom dishes and deprived of amino acids and FBS for 1 h. Total lysosome staining was performed with 1:4000-diluted LysoPrime Green (Dojindo, Kumamoto, Japan) during starvation (1 h), while acidified lysosomes were selectively stained with 100 nM LysoTracker Red DND-99 (Molecular Probes, Eugene, OR) during the latter half of the starvation (30 min). After washout, cells were cultured in phenol red-free D-MEM without amino acids (high glucose and sodium pyruvate; Fujifilm Wako Chemicals), but containing 100 U/ml penicillin G, and 0.1 mg/ml streptomycin under a 5% CO_2_ atmosphere at 37°C, and were then imaged by confocal microscopy.

#### Lipid droplet staining

Cells were grown on glass-bottom dishes and cultured in a medium containing 200 µM oleic acid (Tokyo Chemical Industry, Tokyo, Japan) for 24 h. Lipid droplets were stained with 0.5 µM Lipi-Green (Dojindo) for 30 min. After washout, cells were cultured in phenol red-free D-MEM (high glucose without L-glutamate; Fujifilm Wako Chemicals) containing 8% FBS, 4 mM L-glutamate (Sigma Aldrich), 100 U/ml penicillin G, and 0.1 mg/ ml streptomycin under a 5% CO_2_ atmosphere at 37°C, and were then imaged by confocal microscopy.

#### Nuclei staining

Cells were grown on glass-bottom dishes, treated with appropriate reagents, and simultaneously stained with 20 µM Hoechst 33342 (bisbenzimide; Merck Millipore, Burlington, MA) for 30 min. After washout, cells were cultured in phenol red-free D-MEM (high glucose without L-glutamate; Fujifilm Wako Chemicals) containing 8% FBS, 4 mM L-glutamate (Sigma Aldrich), 100 U/ml penicillin G, and 0.1 mg/ml streptomycin under a 5% CO_2_ atmosphere at 37°C, and were then imaged by confocal microscopy.

### Development of DIC2Cells

#### Implementation and network architecture

The code, installation instructions, and pre-trained model are on GitHub: https://github.com/MitoTAIKI/DIC2Cells-from-OrgaMeas. We provide a Google Colaboratory (Google Colab) notebook, which allows users to test and train DIC2Cells with their own data without having to install the code on their local machine. The network architecture of DIC2Cells uses a conditional adversarial generative network (cGAN) from pix2pixHD, a deep learning technique for high-resolution image-to-image translation, which was implemented on Google Colab in Python 3.7 using the PyTorch machine learning library.

#### Preparation of training dataset

Live cell imaging of HeLa cells stably expressing GFP (clone #1 and #2) was acquired in the form of 8-bit DIC images and corresponding GFP images of 1024 × 1024 pixels. Using ImageJ/Fiji, GT images were generated by manual annotation of individual cells on the GFP images. The DIC and GT images were augmented using the “Create augmented data” algorithm from the MitoS-segmentation-tool available on GitHub. To increase contrast and accentuate details in the images, the DIC images were filtered using the “Sharpen” tool from ImageJ/Fiji.

#### Training of pix2pixHD

The training step was performed using the manually prepared training dataset. The training settings included a load size of 1024 × 1024, batch size of 1, an Adam optimizer, a learning rate of 0.0002, 100 epochs, and a Tesla T4 processing unit from Google Colab PRO. The default loss function in pix2pixHD was monitored during training, and training models were saved every 10 epochs.

#### Validation

The validation dataset consists of DIC and GT images prepared in the same manner as those used for the training dataset (**Table S2**). The 10 DIC2Cells models that were obtained by saving the trained data at every 10 epochs were used to generate segmented cell images (output) from the DIC images in the validation dataset. The Dice coefficient was calculated for the comparison between the output images and corresponding GT images:

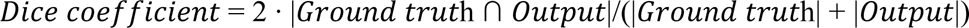

The cell count accuracy (CCA) was also calculated to provide another parameter for evaluating cell segmentation accuracy. In this, true positive (TP) indicates the number of cells that were successfully segmented in comparison with the GT images, whereas false positive (FP) shows the number of cells that were incompletely distinguished from neighboring cells and mistakenly counted as a single cell:

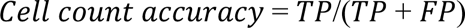

### Using MitoSegNet

The code for MitoSegNet and the pre-trained model were installed on the local machine from GitHub: https://github.com/MitoSegNet/MitoS-segmentation-tool. Live-cell images of mitochondria, lysosomes, lipid droplets, and nuclei were segmented using the pre-trained MitoSegNet model.

### Development of NucSegNet

Using the advanced mode of MitoSegNet, U-Net (a deep learning framework for the segmentation of biomedical images) was trained with nuclear images and corresponding GT images. For nuclei imaging, live cell images of HeLa cells stained with Hoechst 33342 were acquired as 8-bit images at a size of 1024 × 1024. Using ImageJ/Fiji, GT images were generated through the manual annotation of individual nuclei. The nuclei and GT images were augmented using the “Create augmented data” algorithm from the MitoS-segmentation tool (**Table S3**). The training step was performed with the following training settings: crop, 4 tiles; load size, 512 × 512; batch size, 1;, optimizer, Adam; learning rate, 0.00007; and epochs, 20; and an 8GB GPU (GeForce RTX3050) was used. Eighty percent of the dataset was used to train U-Net, while the rest (20%) was reserved for validation of the resulting models (calculation of the Dice coefficient). Monitoring of the loss function and the saving of each model were performed at every epoch. We finally combined the two deep learning tools, MitoSegNet and NucSegNet, into an integrated tool, which we refer to as OrgaSegNet. The code, installation instructions, and pre-trained model of NucSegNet are available on GitHub: https://github.com/MitoTAIKI/NucSegNet-from-OrgaMeas.

## Supplemental material

Fig. S1 shows a comparison of the techniques for labeling organelles. Fig. S2 presents an evaluation of the accuracy of the initially developed 10 DIC2Cells models in the differentiation between neighboring cells in relation to Fig. 2D. Tables S1, S2, and S3 list the settings of data augmentation for development and validation of DIC2Cells and NucSegNet.

## Data availability

ImageJ/Fiji macros for OrgaMeas, and their instructions, are available on GitHub: https://github.com/MitoTAIKI/IJ-macros-from-OrgaMeas.

## Acknowledgments

This work was supported in part by the Platform Project for Supporting Drug Discovery and Life Science Research (Basis for Supporting Innovative Drug Discovery and Life Science Research (BINDS)) from the Japan Agency for Medical Research and Development (AMED) (JP23ama121032), JSPS KAKENHI (grant number JP23KJ1762 to T. Baba, JP21K06069 to S. Tanimura, and JP23K06117 to K. Takeda), the ANRI Fellowship (to T. Baba), and the Shionogi Infectious Disease Research Promotion Foundation (to S. Tanimura). We thank Edanz (https://jp.edanz.com/ac) for editing a draft of this manuscript.

## Author contributions

Conceptualization, Formal analysis, Methodology, Software, Visualization, Writing–original draft: T. Baba; Supervision, K. Takeda; Investigation, Validation; T. Baba, S. Tanimura, A. Inoue; Project administration: T. Baba and K. Takeda; Data curation, Funding acquisition, Resources, Writing–review and editing: all authors.

---

**Figure S1.**
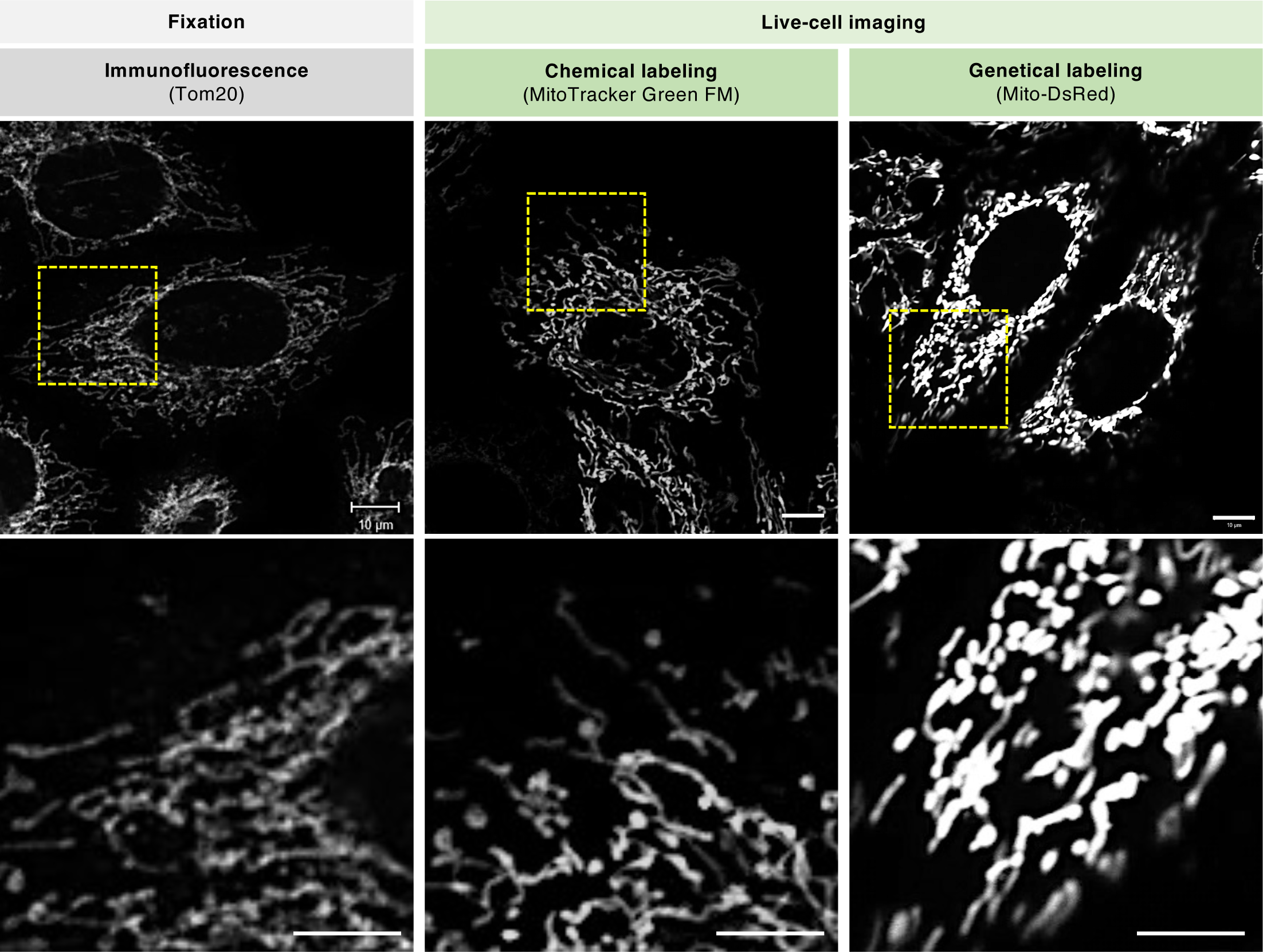
Comparison of the techniques for labeling organelles. HeLa cells were subjected to immunofluorescence with the antibody to the mitochondrial outer membrane protein Tom20 (left panels). Live-cell imaging of HeLa cells labeled with MitoTracker Green FM (middle panels) or transiently expressed with the mitochondrial-targeted fluorescent protein Mito-DsRed (right panels). Scale bar, 10 µm. The lower panels are magnified images of the regions in the yellow dashed lines in the respective upper panels.

**Figure S2.**
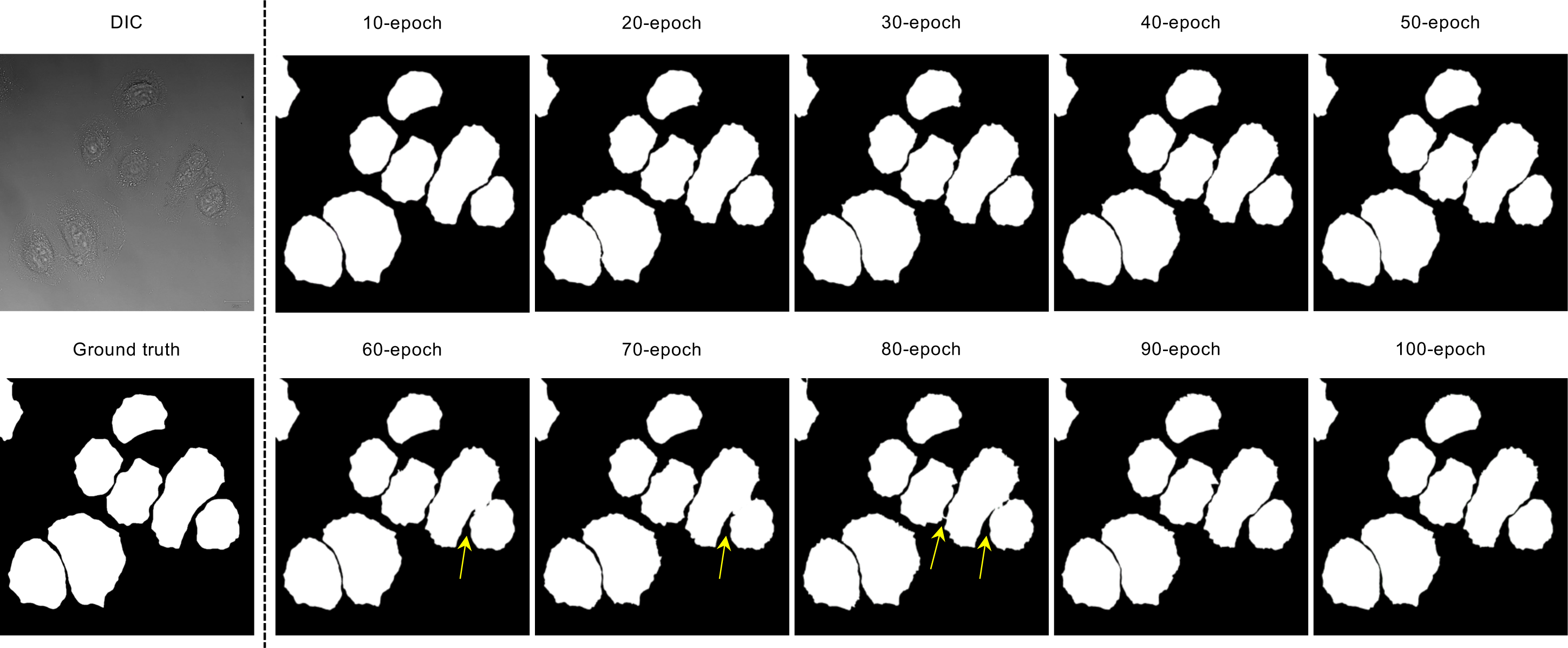
Inconsistencies in the accuracy of the DIC2Cells models in distinguishing between neighboring cells. A representative DIC image (upper left panel), its ground truth image (lower left panel) and output images generated by the 10 DIC2Cells models (other panels) are shown. Yellow arrows indicate the images where the model failed to distinguish between neighboring cells.

**Table S1.** Settings of data augmentation for development of DIC2Cells.

| Data Augmentation |  |
| --- | --- |
| **Horizontal flip** | True |
| **Vertical flip** | True |
| **Width shift range** | 0.3 |
| **Height shift range** | 0.3 |
| **Shear range** | 0.3 |
| **Rotation range** | 180° |
| **Zoom range** | 0.3 |
| **Brightness range** | 0.7 – 1.3 |

**Table S2.**
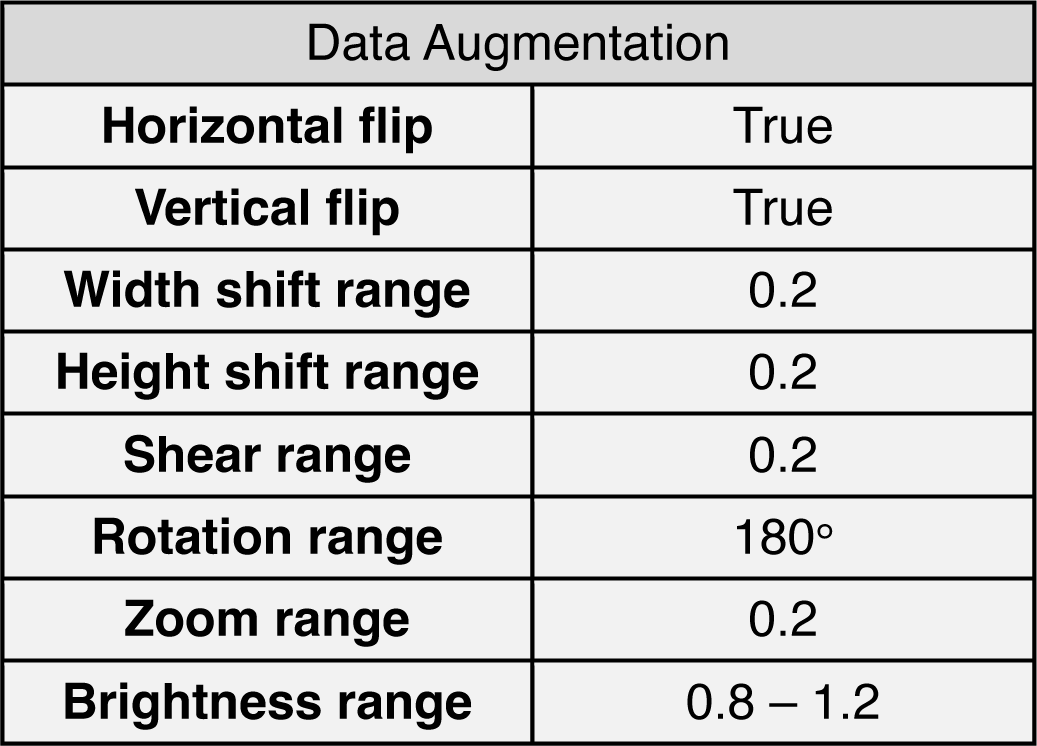
Settings of data augmentation for validation of DIC2Cells.

**Table S3.** Settings of data augmentation for development and validation of NucSegNet.

| Data Augmentation |  |
| --- | --- |
| **Split** | 4 tiles (512 x 512) |
| **Horizontal flip** | True |
| **Vertical flip** | True |
| **Width shift range** | 0.3 |
| **Height shift range** | 0.3 |
| **Shear range** | 0.3 |
| **Rotation range** | 180° |
| **Zoom range** | 0.3 |
| **Brightness range** | 0.7 – 1.3 |

